# Synthetic protein alignments by CCMgen quantify noise in residue-residue contact prediction

**DOI:** 10.1101/344333

**Authors:** Susann Vorberg, Stefan Seemayer, Johannes Söding

## Abstract

Compensatory mutations between protein residues that are in physical contact with each other can manifest themselves as statistical couplings between the corresponding columns in a multiple sequence alignment (MSA) of the protein family. Conversely, high coupling coefficients predict residues contacts. Methods for de-novo protein structure prediction based on this approach are becoming increasingly reliable. Their main limitation is the strong systematic and statistical noise in the estimation of coupling coefficients, which has so far limited their application to very large protein families. While most research has focused on boosting contact prediction quality by adding external information, little progress has been made to improve the statistical procedure at the core. In that regard, our lack of understanding of the sources of noise poses a major obstacle. We have developed CCMgen, the first method for simulating protein evolution by providing full control over the generation of realistic synthetic MSAs with pairwise statistical couplings between residue positions. This procedure requires an exact statistical model that reliably reproduces observed alignment statistics. With CCMpredPy we also provide an implementation of persistent contrastive divergence (PCD), a precise inference technique that enables us to learn the required high-quality statistical models. We demonstrate how CCMgen can facilitate the development and testing of contact prediction methods by analysing the systematic noise contributions from phylogeny and entropy. For that purpose we propose a simple entropy correction (EC) strategy which disentangles the correction for both sources of noise. We find that entropy contributes typically roughly twice as much noise as phylogeny.

## I. INTRODUCTION

In the course of evolution, proteins are under selective pressure to maintain their function and correspondingly their structure. A possible mechanism to maintain structural integrity is the compensation of deleterious mutations between residue pairs in physical contact, known as compensatory mutations: Upon the mutation of one residue the contacting residue has an increased probability to mutate into a residue that will locally restabilize the protein structure, for instance by regaining a lost interaction between them. In multiple sequence alignments (MSAs) of related proteins, this effect leads to correlations between columns of residues in contact among most protein family members [19, 20, 42, 51]. Many of these correlations are indirect, though, and arise through transitive chains of contacting residue pairs [5, 17, 33, 58].

By applying statistical techniques that can distinguish mere correlation from direct statistical coupling of residue positions [33, 54, 58], many false positive predictions could be eliminated. The adoption of this class of statistical models, known as Markov random fields (MRFs), or Potts models in statistical physics, led to a breakthrough in de-novo (template-free) protein structure prediction: The predicted contacts proved sufficiently accurate to be used as spatial restraints to reliably predict protein 3D structures purely from sequence information [23, 26–28, 32, 35, 36, 44–47].

The requirement for large MSAs for sufficiently precise predictions has limited the applicability of contact-assisted de-novo protein structure prediction, all the more because large protein families are more likely to contain at least one member whose structure has been solved and which can be used as a template for homology modelling. Therefore, most research has focused on making contact prediction reliable enough for medium-sized protein families [21, 24, 29, 38, 47, 57].

The background noise effects arising in residue-residue contact prediction have been postulated to arise from three independent sources [1, 8, 14, 18, 22, 33, 37, 43]: random sampling noise due to the limited number of sequences, phylogenetic noise due to the evolutionary relatedness of sequences in the MSA, and entropic noise, which biases high-entropy columns towards higher scores. Unfortunately, the relative contribution and properties of the three different sources of noise are difficult to study in real alignments, mainly because the true values of coupling parameters are not known. In addition, the stochastic noise, entropy-dependent noise and phylogenetic noise cannot be modified independently (for example by sub-sampling), as these noise sources are indirect, complex consequences of learning on only a limited number of sequences that are statistically dependent on each other according to their phylogenetic relationship.

Many correction schemes for removing noise from the matrix of predicted contact scores have been examined [6, 22, 34, 37, 43, 56, 59], and the *average product correction (APC)* [8] came out as a clear winner and is used in almost all recent studies. However, it is widely acknowledged in the field that our limited understanding of what noise effects APC is correcting and why it is so effectively correcting them is severely impeding progress in developing better statistical methods to predict contacting residue pairs. We repeatedly made the experience that a promising extension to the standard MRF model that considerably improved the contact prediction performance *before* applying the APC was doomed to failure because it inexplicably yielded worse results than the baseline method *after* applying APC.

Here, we propose a simple entropy correction (EC) that is computed solely from per-column entropies of the input MSA and corrects for entropy-dependent systematic noise without affecting noise from phylogenetic effects. We find that the EC eliminates nearly as much noise as the APC and therefore suspect that APC also mainly corrects for entropy noise. Whereas the APC is applied as a post-correction to the matrix of predicted contact scores, the EC can be applied directly on the statistical couplings of the MRF model, prior to computing a contact score and other post-processing treatments.

To systematically study the sources of noise limiting the accuracy of contact predictions from MSAs and to facilitate progress in the development of better contact prediction methods, we have developed CCMgen, a method for generating realistic synthetic protein sequence alignments whose residues obey the selection pressures described by a MRF with pairwise statistical couplings between residue positions.

For that purpose, CCMgen requires an exact statistical model that will reliably reproduce the empirical alignment statistics, such as single-site, pairwise or even higher-order amino acid frequencies, of the input MSA that was used to learn the MRF model in the first place. A typical strategy to obtain estimates of the MRF model parameters would involve maximizing the logarithm of the likelihood function over all sequences in the MSA. However, the normalization factor in the likelihood function requires to sum 20*^L^* terms, where *L* is the protein length and methods to optimize the full likelihood are very slow for realistic proteins [4, 12, 33]. The most popular approximation is to maximize the pseudo-likelihood instead of the likelihood, as it can be shown that it converges to the same solution for large numbers of samples and it is fast to compute [2, 10, 50]. Even though pseudo-likelihood maximization gives results of the same quality of predicted residue-residue contacts as those using the full likelihood optimization, several studies unveiled that the pseudo-likelihood model is inaccurate and not able to accurately reproduce the empirical alignment statistics [7, 12].

We provide an implementation of an alternative precise inference technique called persistent contrastive divergence (PCD) [55] with our tool CCMpredPy. Compared to pseudo-likelihood maximization, PCD achieves identical precision for contact prediction while the inferred MRF model reproduces empirical marginals much more precisely. The increased quality of the models comes at the expense of longer run times, which are however still practical for even large proteins and alignments using a single desktop computer. High quality MRF models learned with PCD might prove beneficial beyond the purpose of contact prediction when problems require exact model statistics, e.g. when studying mutational effects or designing new protein features using the model energies.

Finally, we employ CCMgen in combination with MRF models that have been learned with the PCD algorithm and our entropy correction to quantify the relative effect sizes of phylogenetic and entropic noise on the precision of contact prediction. We find that the contribution of entropy noise in contact prediction is on average twice as big as that of phylogenetic noise.

## II. RESULTS

### Persistent Contrastive Divergence Allows Accurate Inference of MRFs

An exactly inferred MRF will reliably reproduce the empirical single-site and pairwise amino acid frequencies, *f_i_*(*a*) and *f_ij_*(*a, b*) for all positions *i, j* in the MSA and all amino acids *a, b* ∈ {1, …, 20} [40, 58]. Several studies demonstrated that pseudo-likelihood maximization, while being the method of choice for contact prediction, yields models that cannot accurately reproduce the empirical alignment statistics [4, 7, 12].

We developed a method that uses an inference technique called persistent contrastive divergence (PCD) [55] to learn MRF models that accurately reproduce the empirical alignment statistics. As in the study by Figliuzzi *et al*. [12], we computed for all Pfam MSAs in the PSICOV dataset the single-column and paired-column amino acid frequencies as well as covariances, *cov*(*δ_a,xi_, δ_b,xj_*) = *f_ij_*(*a, b*) – *f_i_*(*a*) *f_j_*(*b*), where *δ_a,x_* is the Kronecker symbol. We compared these statistics with those from sequences obtained by MCMC sampling from MRFs that were trained on the Pfam MSAs using either pseudo-likelihood maximization or PCD.

We find that the empirical single-site amino acid frequencies are well reproduced by both models. But whereas the empirical pairwise amino acid frequencies and covariances correlate strongly with the corresponding statistics computed from the PCD samples, the correlation is much weaker for samples obtained from pseudo-likelihood MRF models (Figure 1a and 1b and Supplemental Figure 1).

**FIG. 1.**
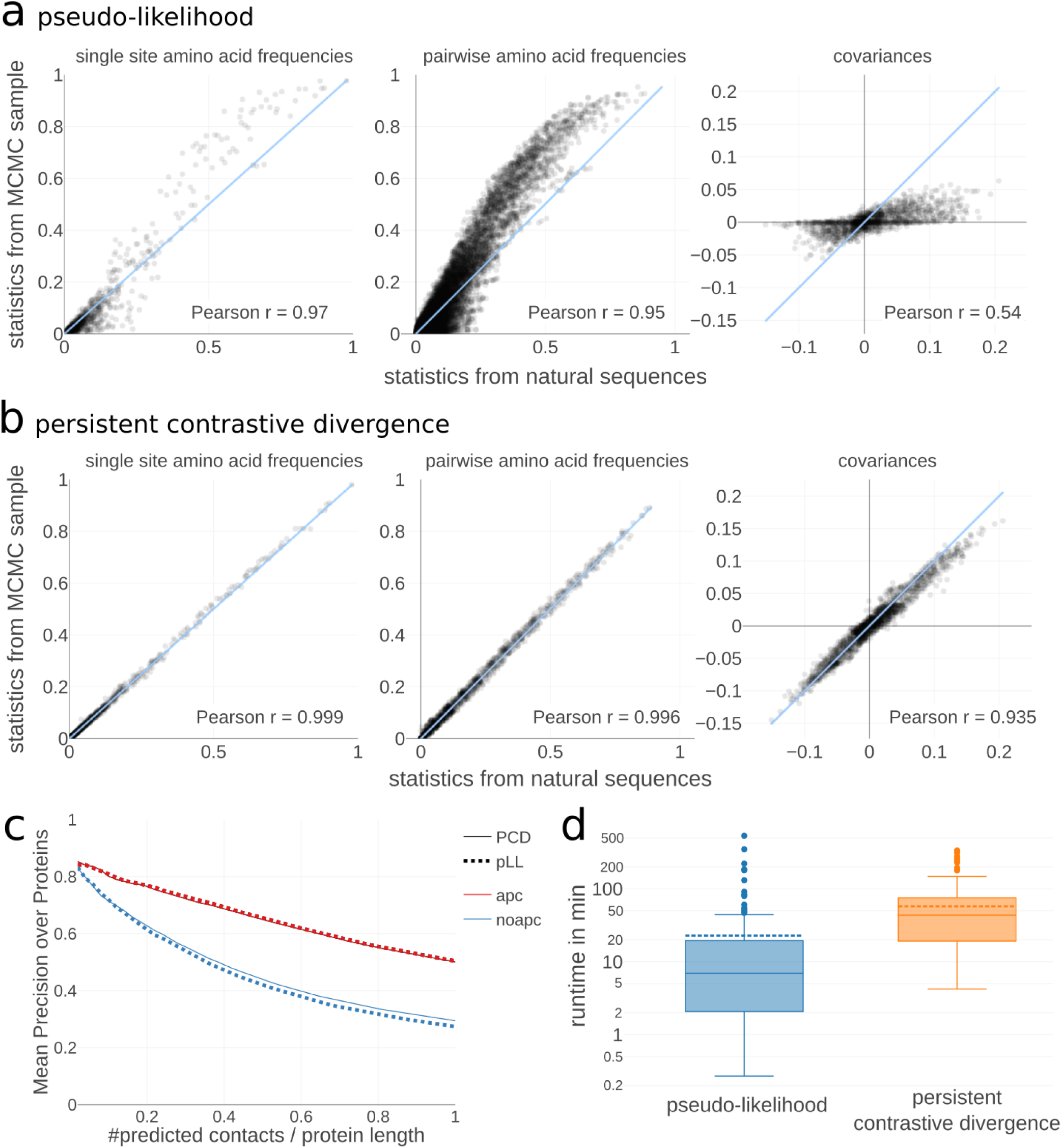
Persistent contrastive divergence permits the inference of high-quality models. **a** and **b** Comparing single amino acid frequencies (left), pairwise amino acid frequencies (center) and covariances (right) computed from natural sequences (Pfam alignment) and from Markov chain Monte Carlo (MCMC) samples generated from Markov random field (MRF) models trained with either pseudo-likelihood maximization (**a**) or persistent contrastive divergence (PCD) (**b**) for protein 1bkrA in the PSICOV dataset. **c** Mean precision of contacts predicted as (APC corrected) *L*_2_ norm of pair couplings of a MRF trained with either pseudo-likelihood maximization or PCD. **d** Distribution of run times in minutes for the 150 proteins in the PSICOV dataset when learning MRF models with CCMpredPy (run on 4 cores). The median runtime in minutes with pseudo-likelihood is 7 minutes and with persistent contrastive divergence is 43.5 minutes. Dashed line in boxplots represents the mean, solid line represents the median of the distribution.

Furthermore, as in Figliuzzi *et al*. [12], we investigated how well the generated MCMC samples reproduce the alignment substructure of the original Pfam alignments with respect to the organisation of subfamilies in sequence space. We projected the protein sequences of the MCMC samples onto the first two principle components obtained from a principal component analysis (PCA) of the original Pfam MSA. Again, we find that the alignment substructure described by the grouping of sequences that can be observed in the two-dimensional PCA space, is reproduced more reliably by MCMC samples generated from PCD models than from pseudo-likelihood models (Supplemental Figure 2b and 2d).

It has been argued that for the purpose of predicting residue contacts an approximate model such as those obtained by maximizing the pseudo-likelihood for a MRF is sufficiently accurate to infer the correct topology of the interaction network of residues [7]. Indeed, predicted contacts from a PCD model achieve equal precision as predictions from a pseudo-likelihood model (Figure 1a). In the Supplemental information, further analysis is shown comparing the APC-corrected contact scores from pseudo-likelihood and PCD models (Supplemental Figure 3).

However, more complex problems such as prediction of mutational effects or generating realistic samples of sequences, require exact model statistics. Several methods have been developed that exactly infer MRF models, such as ACE [4], bmDCA [12] or general MCMC methods [33], but they are computationally intensive which renders them impractical for real proteins. In comparison, our PCD-based CCMpredPy method is only about a magnitude slower than pseudo-likelihood maximization (Figure 1b).

### Correcting for Entropy Bias Removes a Major Source of Noise

A major obstacle for improving the statistical methods for residue-residue contact prediction is our lack of understanding of the sources of noise. The background noise effects have been postulated to arise from at least three independent sources, whose size and properties are difficult to quantify.

*Phylogenetic noise* originates from the violation of the assumption of independence of sequences in the MSA. This assumption has been made by all methods that have been employed for contact prediction so far.

To understand the origin of phylogenetic noise, consider the example in Figure 2. The MSA is composed of two subtrees whose last common ancestor sequences, DSMF and ETMF, had a mutation at the second and first position respectively. All descendants of the first ancestral sequence whose first two residues have not mutated in the meantime will have a DS at first and second position, while all descendants of the other ancestral sequence whose first two residues have not mutated yet will have a ET at those positions. Therefore, pairs DS and ET are more likely than would be expected from the frequencies of D and E in the first column and of S and T in the second column. The first and second position will therefore appear to be statistically coupled even though they are not.

**FIG. 2.**
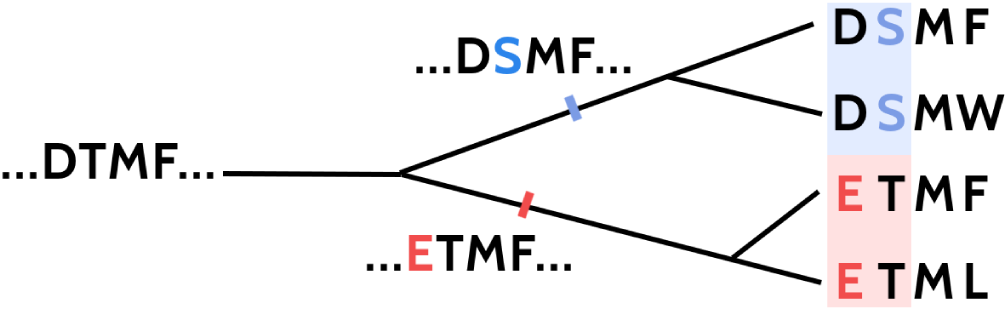
The phylogenetic dependence of closely related sequences can produce covariation signals. Here, two independent mutation events (highlighted in red and blue) in two branches of the tree result in a covariation signal for the two affected positions.

*Entropic bias* leads to higher contact scores the higher the entropies of columns *i* and *j* are. To understand the reason behind this, let us assume that a MSA has no statistically coupled residue pairs. For two relatively well conserved positions *i* and *j*, where column entropies are low, many of the 400 amino acid pairs (a, b) will not be observed. The commonly required regularization of the MRF will ensure that those amino acid pairs (*a*, *b*) without counts will not contribute to the overall contact score *c_ij_* (e.g. as described in equation 5) for this residue pair. The higher the entropies of columns *i* and *j* are, the more of the 400 amino acid pairs will receive some of the *N* total counts. The fraction of observed (*a*, *b*) pairs, *f_ij_*(*a*, *b*), will only rarely be *exactly* equal to *f_i_*(*a*) × *f_j_*(*b*), the fraction of sequences with *x_i_* = *a* times the fraction with *x_j_* = *b*. Each such pair (*a*, *b*) will therefore make a small contribution to the overall contact score *c_ij_*. The more amino acid pairs (*a*, *b*) in columns *i* and *j* receive at least one or a few counts, the higher will be the expectation value of the contact score *c_ij_* by chance in the absence of any coupling.

*Sampling noise* on the estimated coupling coefficients would remain, even if we were able to correct for entropic bias and phylogenetic effects, because with a finite sample of sequences we cannot estimate fractions arbitrarily accurately. For example even if the sequences could be assumed to be independent of each other, the probability of an amino acid pair (*a, b*) that has been observed *n* ≪ *N* times out of *N* is only estimated to a relative accuracy of approximately 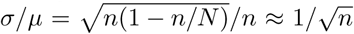, according to the standard deviation of the binomial distribution.

Figure 3a shows the contact scores in grey scale computed from a MRF that has been trained from a typical example MSA (for details on how to obtain contact scores see Materials and Methods). The striping patterns in horizontal and vertical directions reflect strong systematic row- and column-dependent score biases. Some positions seem to obtain generally higher scores than others. Without correction, ranking by these scores would severely overpredict contacts between positions with positive score bias and underpredict contacts between positions with negative bias.

**FIG. 3.**
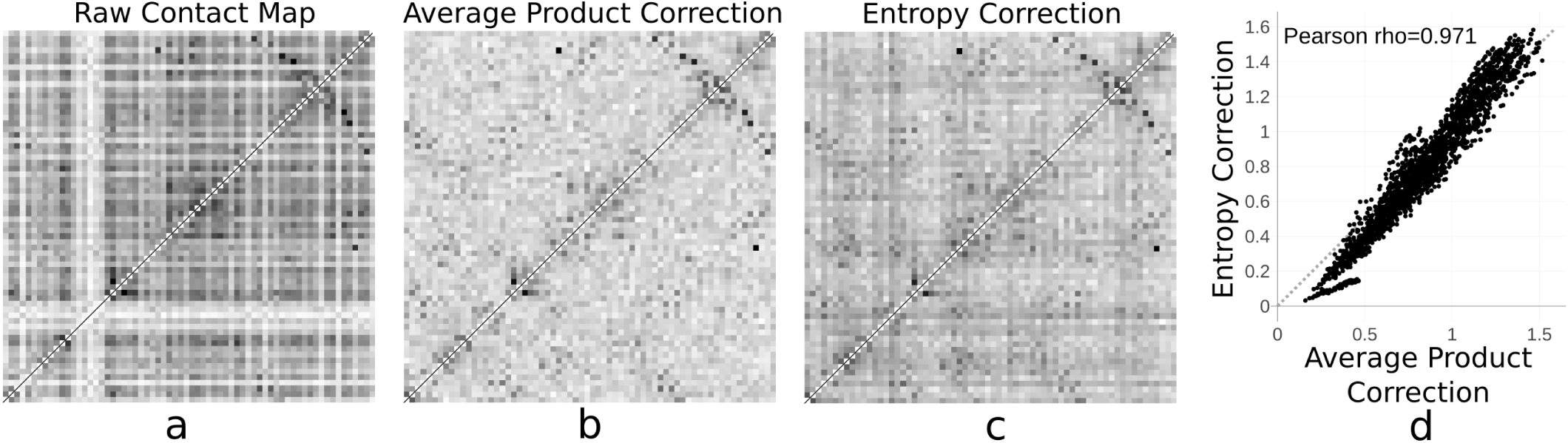
Entropy correction eliminates major source of noise. Raw and corrected contact score matrices for protein 1c9oA. The gray scale indicates the contact scores for each residue pair (*i*, *j*) for raw uncorrected scores computed from a MRF model trained with PCD (**a**, equation 5
), average product corrected scores (**b**, equation 6) and entropy corrected scores (c, equation 7). The striping pattern in **a** arises from systematic score biases, which originate mainly from entropy. **d** For protein 1c9oA, the correction term defined by APC correlates well (Pearson *ρ* = 0.971) with the correction determined by our EC strategy.

Applying the APC eliminates the systematic effects leading to the striping patterns (Figure 3b). It thereby greatly improves the performance of all contact prediction scores for local pairwise measures such as mutual information as well as for global statistical coupling methods such as the MRF-based score referred to here [5, 8, 9, 30, 31].

To disentangle the entropic noise from the phylogenetic bias, we first propose an entropy-dependent correction, EC, of the contact scores *c_ij_* (see equation 5) that depends solely on the per-column entropies *s_i_* of the MSA from which the MRF was trained,
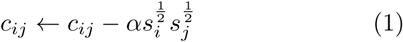

with an analytically determined constant *α* that depends on all column entropies *s*_1_, …, *s_L_* (for details see Materials and Methods).

Figure 3c shows how the entropy correction removes almost as much of the striping effects as APC and Figure 3d reveals how strongly both corrections correlate (see Supplemental Figure 4 for the whole data set). Apparently, APC mainly corrects out entropy bias [8].

However, the relative contributions of entropic and phylogenetic noise limiting the precision of contact prediction are yet unclear. In the following we will use our tool CCMgen to distinguish between both sources of noise.

### Quantifying Noise Effects with CCMgen Reveals Entropy as Dominating Source of Noise

Our workflow to analyse the relative contributions of noise sources is described in Figure 4. First, we estimate the parameters of a second order MRF model with PCD using CCMpredPy for each of the 150 Pfam MSAs in the PSICOV data set.

**FIG. 4.**
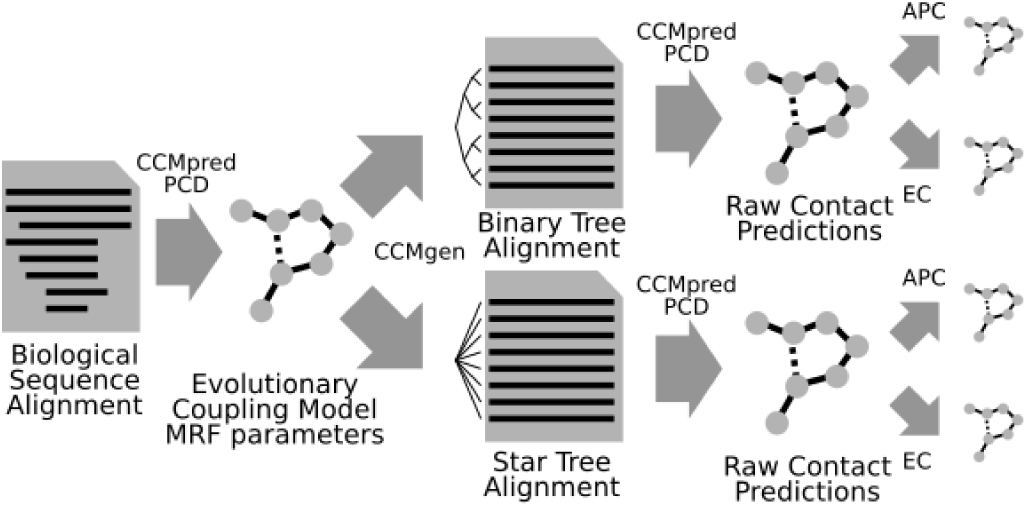
Workflow for quantifying noise effects. Artificial multiple sequence alignments are generated using a binary tree phylogeny for sequences with strong interdependencies and using a star tree phylogeny for nearly independently sampled sequences and then contacts are predicted and post-corrected for both sets of alignments.

In a second step, we use CCMgen with the learned model parameters to generate realistic synthetic MSAs of interdependent sequences with pairwise statistical couplings between some positions as they are observed in MSAs between residues in physical contact. CCMgen provides full control over the generation of the synthetic MSAs by allowing us to specify the evolutionary times and phylogeny along which the sequences are sampled. We sample two sets of synthetic MSAs: one set with a star tree topology and the other with a binary tree topology (Figure 5). Given sufficient evolutionary time, the phylogenetic dependencies between sequences drawn according to the star tree topology should be negligible, whereas sequences drawn along the binary tree are expected to contain stronger interdependencies.

**FIG. 5.**
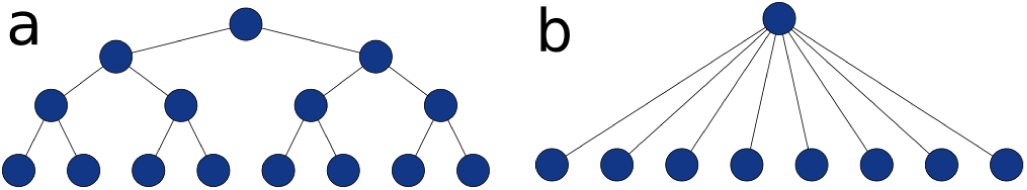
CCMgen can generate MSAs based on a MRF model and a phylogenetic tree supplied either as Newick file or as one of the two shown, idealized topologies: **a** binary tree and **b** star-shaped tree.

Because the accuracy of predictions strongly depends on alignment depth and diversity [30, 39], we ensured that the synthetic alignments contain the same number of sequences and have similar diversities as the original Pfam alignments (for details see Material and Methods). These provisions justify a direct comparison of the results for sampling sequences along the star and binary topologies.

Third, we run CCMpredPy on each of the synthetic MSAs and predict residue-residue contacts by ranking the pairs according to the descending raw coupling scores (equation 5), or by the APC-corrected coupling scores (equation 6) or by entropy corrected scores (equation 7). Since we know the ground truth of which pairs are coupled from the MRF model used for generating the synthetic MSAs, we can use these alignments to investigate and quantify the effect of phylogenetic noise on the precision of residue-residue contact prediction.

Figure 6a and 6b plot the mean precision of the predicted contacts versus the number of predictions per columns in the MSA. The precision for one MSA is the fraction of correctly predicted contacting pairs of positions (*i, j*) out of all predicted pairs. The correctly predicted pairs (*i, j*) are those for which the *C_β_*−*C_β_* distance in the reference protein structure of the Pfam MSA is below 8Å. As expected, the mean precision drops as more predictions are considered and lower ranks are included.

**FIG. 6.**
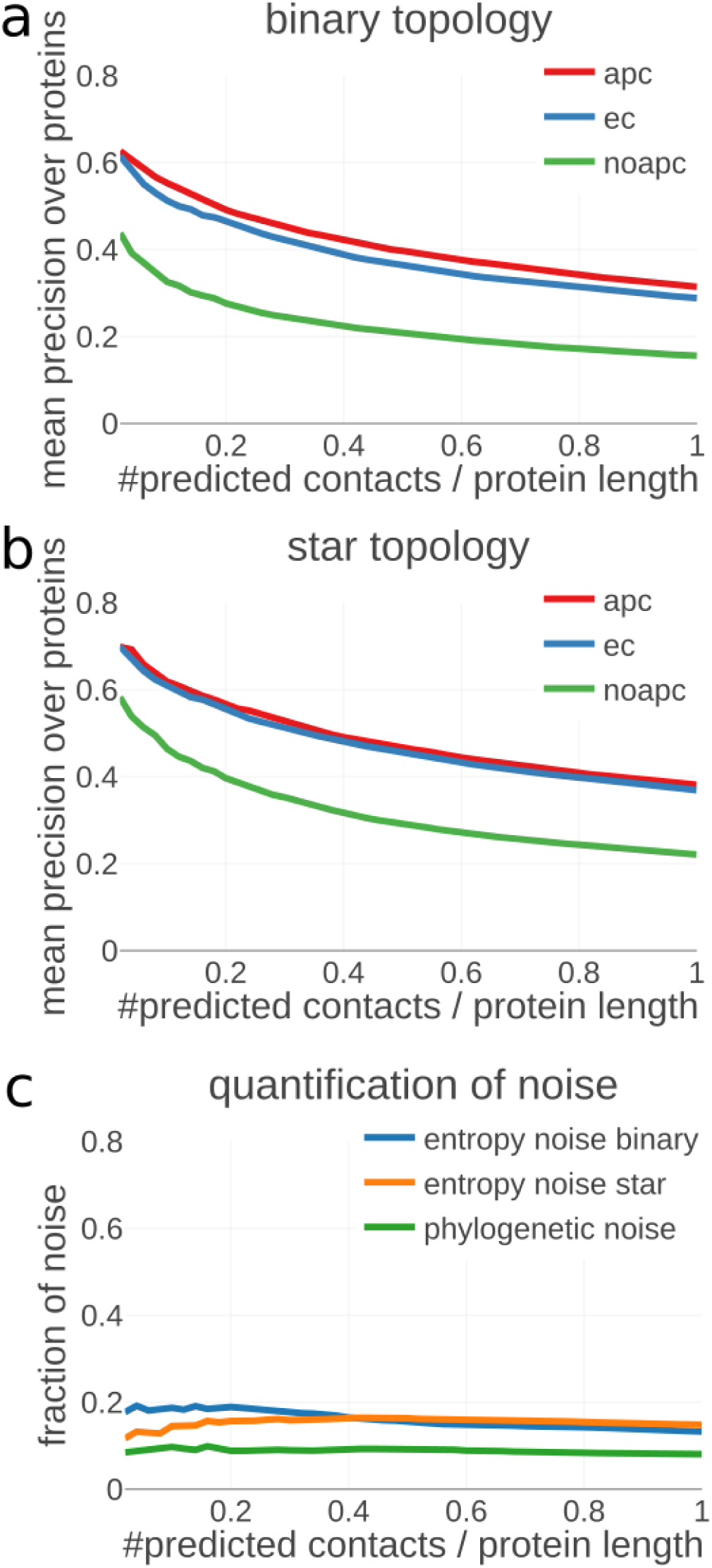
Effect of phylogenetic noise on contact prediction accuracy. Since the average product correction (APC) corrects for both phylogenetic and entropic effects and the entropy correction (EC) corrects only for entropic effects, they exhibit similar contact precision profiles when sequences are drawn according to a star tree topology (**a**) but the differences in phylogenetic and entropic noise become apparent when sequences are phylogenetically coupled (**b**). Since star trees do not have phylogenetic noise, we can estimate the contributions of phylogenetic and entropic noise, which are about one-to-two in strength (**c**).

Both APC and EC correction have a huge effect in reducing noise and increasing the precision of predictions. While the EC can only correct entropy-dependent noise, the APC should also partially correct phylogenetic noise, because this noise source affects some positions more than others (Figure 2). Indeed, we observe that both corrections give very similar results for MSAs generated with star topology trees, which are not expected to show phylogenetic noise, while the APC outperforms the EC on MSAs with binary tree topologies. We therefore interpret the performance gap between the APC and EC on binary trees as phylogenetic noise at least partially corrected by the APC but not by the EC.

We estimate the strength of the phylogenetic noise as the drop in precision between the EC-corrected precisions on MSAs with star topology and EC-corrected precision on the MSAs with binary tree topology (Figure 6c). The strength of the entropy noise is shown in terms of the drop in precision between the EC-corrected and uncorrected, raw coupling scores, both for the star tree topology and for the binary tree topology. The contribution of entropy noise to the drop in precision is roughly two times larger than that of the phylogenetic noise.

## III. DISCUSSION

### Persistent Contrastive Divergence Facilitates Inference of High Quality Models

Pseudo-likelihood maximization is the state-of-the-art inference technique for MRF models in contact prediction. Whereas the approximate nature of the model is sufficient for the correct ranking of residue pairs, the model is not exact in a way that it can reliably reproduce the empirical amino acid statistics of the original MSA. We implemented an alternative inference technique for MRFs, known as persistent contrastive divergence (PCD) which yields similar precision for predicted contacts but permits learning the fine statistics of the MRF model with higher precision. Even though other accurate model inference methods such as ACE [4] or bmDCA [12] can infer model parameters up to arbitrary precision, they are computationally intensive and their applicability is limited to small proteins. Our open source Python implementation of the PCD algorithm which is available through our tool CCMpredPy, is only about 7 times slower than pseudo-likelihood maximization. Hence it might be of use for large-scale studies that require exact models, such as investigating mutational effects or designing new protein features.

### Entropy Correction Eliminates Major Source of Noise

The average product correction (APC) and other de-noising techniques such as LRS [59] estimate a background model from the raw contact score matrix, which therefore depends on the statistical model used to infer the contact matrix. This means that when attempting to improve contact prediction through an altered probabilistic model, an equivalent procedure to the APC has to be found for the new method to be competitive. Our definition of an entropy correction (EC) is computed from statistics of the input MSA and has the valuable property of factorizing into a term depending only on *i* and one depending only on *j*.

In fact, we derived the here proposed EC when calculating the expectation value correction of the contact score based on the null model assumption of no couplings in which the pairwise amino acid counts should follow a hypergeometric distribution. The expectation value correction for squared couplings *w_ij_*(*a*, *b*)^2^ can be formulated as
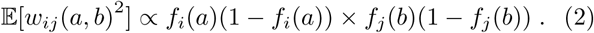

This equation allows for the correction of individual couplings *w_ij_*(*a*, *b*). It could therefore be used to train deep neural networks directly on the EC-corrected coupling coefficients *w_ij_*(*a*, *b*), combining the advantages of entropy correction with learning directly from the full set of coupling coefficients [21, 29] instead of only from their EC-corrected norms ∥*w_ij_*∥, as given in equation 5.

Surprisingly however, the previous correction gave slightly worse results than the closely related entropy formulation, which is why we used it in this study. We believe that replacing the APC as a post-hoc correction by a well-understood and statistically motivated correction that can be applied directly on the coupling coefficients removes a major roadblock impeding the progress in developing improved statistical methods. Furthermore, by using our EC strategy it becomes possible to disentangle the correction of entropic effects from correcting for other effects, thereby facilitating the development of improved techniques for directly modelling both entropic and phylogenetic noise and making contact prediction widely applicable.

### CCMgen Allows the Generation of Realistic Synthetic Alignments

We developed CCMgen, the first tool for generating realistic MSAs of protein sequences for a given phylogenetic tree whose residues follow the pairwise coupling constraints from a Markov random field model. CCMgen provides full control of parameters that determine the interdependencies between sequences through the specification of the phylogenetic topology and the evolutionary rate of the sampling process. It enables to distinguish different sources of noise observed in alignments and how they affect the performance of residue-residue contact predictions. We believe CCMgen will prove to be useful for improving and validating contact prediction methods.

In this study, we demonstrated how CCMgen can be applied to analyse the noise contributions from entropy and phylogeny. Given MRF models learnt on real MSAs, we generated synthetic MSAs with statistically coupled amino acid columns from two types of phylogenetic trees, one in which the sequences are maximally independent (star topology) and one in which the statistical dependences are much stronger (binary tree). By predicting contacts from the two types of synthetic alignments and correcting the predicted contacts either with the APC or with our proposed entropy correction, we were able to elucidate the effect of phylogenetic and entropic noise on contact prediction accuracy.

According to the quantification of noise effects, the most important goal for residue-residue contact prediction is an accurate treatment of entropic effects, as they account for roughly two times the amount of correctable noise and are especially important for correctly identifying the strongest evolutionary couplings. However, phylogenetic noise has an important contribution to the predictions and only a fraction of it is probably corrected by the popular average product correction (APC). This result shows that it might be very worthwhile to develop methods for contact prediction and for learning of MRFs that can explicitly take the statistical dependencies of sequences by common descent into account.

## IV. MATERIALS AND METHODS

### Recap: MRFs model statistical couplings between columns in a MSA

To predict contacts between residues, a popular approach is to train a Markov random field (MRF) model describing the probability to observe a sequence **x** = (*x*_1_, …, *x_L_*) of *L* amino acids,
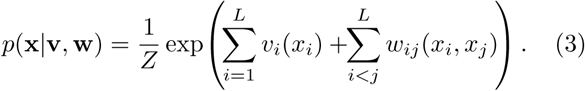

The couplings *w_ij_*(*a, b*) describe the preference to find amino acid *a* at position *i* and *b* at *j* in the same sequence in relation to the probability if these positions were independent, as parametrized by the single-column amino acid preferences *v_i_*(*a*). *Z* is the normalization constant, equal to the sum of the exp function in the numerator summed over all possible 20*^L^* sequences.

To estimate the parameters *v_i_*(*a*) and *w_ij_*(*a*, *b*) of the MRF, the logarithm of the likelihood for all sequences in the MSA, equal to the sum over the log-likelihood of each sequence, could be maximized. A regularization term that pushes all parameters towards zero needs to be added to prevent overtraining, most commonly a *L*_2_ penalty,
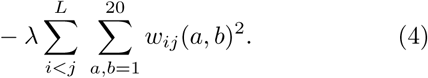

But the huge number 20*^L^* of terms in *Z* renders an exact solution infeasible for realistic protein lengths.

A number of approximations have been developed for this general class of problems. The approach that has consistently been found to work best for residue contact prediction is the pseudo-likelihood approximation, in which we replace the likelihood with the pseudo-likelihood and maximize the regularized log pseudo-likelihood [2, 10, 50],

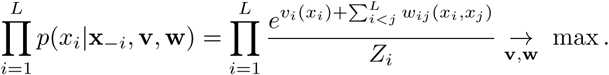

Here, **x**_−*i*_ denotes the vector obtained from **x** by removing the i’th component. The normalization constant *Z_i_* is the sum of the exp function in the numerator over only 20 terms (*x_i_* = 1, …, 20) and can therefore be evaluated easily.

Once the parameters **v, w** are estimated from a MSA, we can predict contacts for pairs of positions *i* and *j* using their statistical couplings. The most often used score for residue contact prediction simply takes the *L*_2_ norm of the 20 × 20-dimensional vector **w***_ij_* with components *w_ij_*(*a*, *b*) [3, 10, 11, 31, 50],
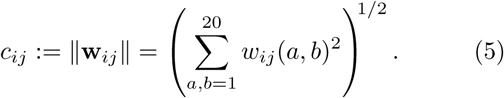

### Recap: Average Product Correction

The APC subtracts from each score *c_ij_* the product of the average score *c_i_*_•_ for row *i* times the average score *c_j_*_•_ for column *j* divided by the average score *c*_••_ over all cells [8]:
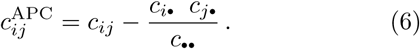

The APC ensures that the average of the corrected coupling score over each column and over each row is 0. This can be verified by summing equation (6) over all *i* or *j*. The assumption made is that, since each residue is only in contact with a small fraction of all residues, the mean coupling score over a column or row is dominated by the systematic score bias on all pairs in the column or row rather than by the coupling scores on a small fraction of contacting residues. APC can also be interpreted as an approximation to the first principal component of the raw contact matrix [59]. It therefore removes the highest variability in the raw contact matrix that is assumed to arise from background biases.

### Entropy Correction

We define an entropy correction (EC) that depends solely on the per-column entropies of the MSA from which the MRF was trained:
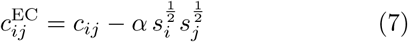

where
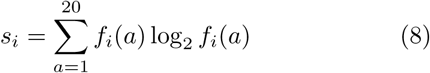

is the entropy of column *i* and *α* is a coefficient determining the strength of the correction. We determine *α* by analytically minimizing the sum of squares of the corrected off-diagonal coupling scores,
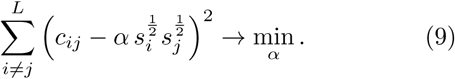

By setting the derivative to zero we obtain the optimal *α* value,
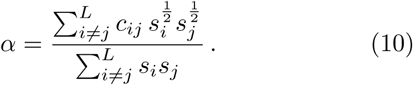

We also investigated other correction strategies using entropy statistics computed from the input MSA, such as the joint entropy for pairs of columns or different exponents in equation 8. The resulting variations of the entropy correction performed comparably regarding the average correlation with APC as well as precision of contact predictions (data not shown).

### Learning MRFs with Persistent Contrastive Divergence

While the log likelihood function cannot be efficiently computed because of the exponential complexity of the normalization constant *Z*, it is possible to approximate its gradient with an approach called contrastive divergence [25].

The gradient of the log likelihood with respect to the couplings *w_ij_*(*a*, *b*) can be written as:
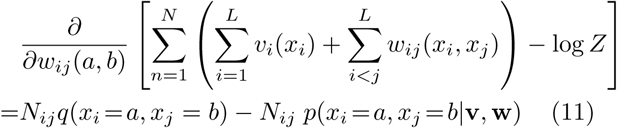

where *N_ij_* represents the number of sequences that do not contain a gap at positions *i* and *j*, *q*(*x_i_* = *a*, *x_j_* = *b*) represents the empirically observed pairwise amino acid frequencies that are normalized over *a, b* ∈ {1, …, 20} and *p*(*x_i_* = *a*, *x_j_* = *b*|**v, w**) corresponds to the model probabilities of the MRF for observing an amino acid pair (*a*, *b*) at positions *i* and *j*. The empirical amino acid counts, given by *N_ij_q*(*x_i_* = *a*, *x_j_* = *b*), are constant and need to be computed only once from the alignment.

The marginal distributions of the MRF cannot be computed analytically as it involves the normalization constant *Z*. Markov chain Monte Carlo (MCMC) algorithms can be used to generate samples from probability distributions that involve the computation of complex integrals such as the normalization constant *Z*. Given that the Markov chains run long enough, the equilibrium statistics of the samples will be identical to the true probability distribution statistics. Thus, an estimate of the marginal distribution of the MRF in the gradient in equation 11, *p*(*x_i_* = *a*, *x_j_* = *b*|**v**, **w**), can be obtained by simply computing the expected amino acid counts from MCMC samples. However, MCMC methods require many sampling steps to obtain unbiased estimates from the stationary distribution which comes at high computational costs.

Hinton suggested contrastive divergence (CD) as an approximation to MCMC methods [25]. The idea is simple: instead of starting a Markov chain from a random point and running it until it has reached the stationary distribution, we run *C* chains in parallel, each being initialized with one of the sequences from the input MSA and we evolve them for only a small number of steps. Obviously the chains do not converge to the stationary distribution in only a few steps and the sequence samples obtained from the current configuration of the chains present biased estimates. The intuition behind CD is that even though the resulting gradient estimate from the biased samples will also be noisy and biased, it points roughly into the same direction as the true gradient of the full likelihood. Therefore the approximate CD gradient should become zero approximately where the true gradient of the likelihood becomes zero.

We apply CD and generate sequence samples to estimate the marginal probabilities by evolving Markov chains which have been initialized with randomly selected protein sequences from the original Pfam MSAs for one full step of Gibbs sampling. We set the number of Markov chains to *C* = max (500, 0.1*N*), with *N* being the number of sequences in the MSA, which seems to give a good trade-off between performance and runtime. Gibbs sampling requires updating each sequence position *x_i_* with *i* ∈ {1, …, *L*} for sequences of length *L* by selecting a new amino acid *a* for that position according to the conditional probability

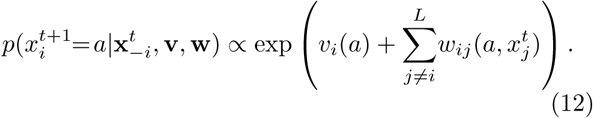

This Gibbs sampling approach is known to generate samples **x**^0^, …, **x***^t^* that are distributed according to the model probability in equation 3 [15, 16]. Note that we do not update positions representing a gap and we thereby retain the gap structure of the initial sequence.

A modification of CD known as persistent contrastive divergence (PCD) does not reinitialize the Markov chains at data samples every time a new gradient is computed [55]. Instead, the Markov chains are kept persistent: they are evolved between successive gradient computations. The assumption behind PCD is that the model changes only slowly between parameter updates given a sufficiently small learning rate. Consequently, the Markov chains will not be pushed too far from equilibrium after each update but rather stay close to the stationary distribution [13, 41, 55].

Tieleman and others observed that PCD performs better than CD in all practical cases tested, even though CD can be faster in the early stages of learning [41, 53, 55].

Therefore we start optimizing the full likelihood with CD and switch to PCD at later stages of learning. CCMpredPy settings for training a MRF with persistent contrastive divergence are listed in Supplementary Materials.

### Sampling Sequences from Phylogenies Using MRFs with CCMgen

Instead of evolving sequences along a linear path, the MRF model can also be used to sample protein sequences according to an arbitrary phylogenetic tree.

CCMgen can simulate the evolution of sequences along any given phylogenetic tree constrained by a MRF model, such as those calculated from CCMpred for example. The user can either supply a phylogenetic tree in Newick format that has been generated by a phylogenetic reconstruction program such as FastTree [48] on a real alignment or choose between two types of idealized trees, a binary and a star-shaped topology. For these idealized trees the user can specify the number of leaf nodes and the total depth of the tree, which is the total number of mutations per position from the sequence at the root to the leaf nodes. The root sequence can either be supplied by the user or be generated by evolving an all-alanine sequence with a number of mutations (i.e. Gibbs sampling steps according to the MRF as described in equation 12). Sequences at subsequent child nodes are generated one by one, by duplicating the sequence at the parent node and evolving the respective child node sequences each with a number of mutations proportional to the edge length. The output of CCMgen is a MSA file with the sequences at the leaf nodes of the tree. CCMgen is released as open-source python command-line application.

### Generating MCMC samples from MRFs with CCMgen

MCMC samples have been generated with CCMgen by evolving 10000 Markov chains by repeated Gibbs sampling as described in equation 12. The Markov chains, each representing protein sequences of length *L* (length of protein in the PSICOV data set) have been randomly initialized with the 20 amino acids. Since the alignment substructure is strongly impacted by the non-random distribution of gaps in the sequences (Supplemental Figure 2a), in a second step the gap structure of randomly selected sequences from the original Pfam alignment is copied over (gaps represented as 21st amino acid). Thus, it is ensured that the sampling procedure reproduces the original alignment substructure as closely as possible (Supplemental Figure 2b and 2c). The number of Gibbs steps before drawing samples was set to 500. Increasing the number of Gibbs steps to e.g. 1000 does not change the statistics of the MCMC samples, hence we can assume that the Markov chains have reached the equilibrium distribution. CCMgen settings for MCMC sampling are listed in Supplementary Materials.

### Workflow for Quantification of Noise in Contact Prediction with CCMgen

We used CCMpredPy to learn MRF models for all Pfam alignments in the PSICOV data set using PCD. In order to obtain models with few but precise constraints, we fixed gradients of coupling parameters at zero for non-contacting residue pairs (*C_β_* distance > 12Å) during training. This procedure ensures that the majority of residue pairs not forming contacts in the protein structure will not be coupled in the MRF model.

We used CCMgen with the learned MRF models to generate synthetic alignments by evolving sequences along idealized star and binary tree topologies. The ancestral sequence at the root of a tree, **x***^t^* with *t* = 0, is obtained by evolving an all-alanine sequence for 10 steps of Gibbs sampling as described in equation 12. The synthetic alignments have the same number of sequences as the corresponding Pfam alignments from the PSICOV set.

We also ensure that the diversity of the resulting MSAs is within 1% of the diversity of the original Pfam MSA from the PSICOV set by adjusting the depth of the trees, which is equivalent to adapting the mutation rate (Supplemental Figure 5a). We measure diversity as the number of effective sequences, *N*_eff_, defined as the exponential of the average column entropies, as defined in the HH-suite software package [49].

For each new CCMgen run with adapted mutation rate a new ancestral sequence is sampled by evolving an all-alanine sequence for 10 Gibbs steps. We found that the aforementioned procedure of masking non-contacting residue pairs during training of the MRF model has the advantageous effect of allowing smaller mutation rates to be used to achieve the desired diversity compared to sampling sequences with a fully parametrized model, likely due to the smaller number of constraints trapping the sampling procedure in local optima. Enforcing small mutation rates is essential for preserving the interdependence between sequences when sampling along binary topologies, which consequently controls the amount of phylogenetic bias. The mutation rates used to obtain synthetic alignments of similar diversity as the original Pfam alignments are very similar regardless of the phy logenetic topology along which sequences were sampled (Supplemental Figure 5b).

CCMpredPy and CCMgen settings for training the MRF with persistent contrastive divergence and generating the synthetic alignments along binary and star tree topologies are listed in Supplementary Materials.

### Dataset

We used the PSICOV data set that was published together with the PSICOV method [30] and which comprises MSAs for 150 Pfam domains with known crystal structures. For each Pfam MSA in the PSICOV set we first removed sequences with more than 75% gaps and columns with more than 50% gaps, similarly as in Ekeberg *et al*. [9], Kamisetty *et al*. [31], Skwark *et al*. [52], to reduce the well-known impact of gaps on the analysis.

Sequences in a MSA do not represent independent draws from a probabilistic model. To reduce the effects of redundant sequences, we employ a popular sequence reweighting strategy that has been found to improve contact prediction performance. Every sequence *x_n_* of length *L* with *n* ∈ {1, …, *N*} in an alignment with *N* sequences has an associated weight *w_n_* = 1/*m_n_*, where *m_n_* represents the number of similar sequences:
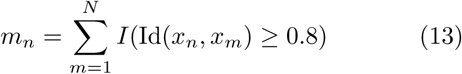

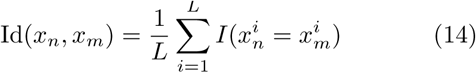

An identity threshold of 0.8 has been used for all analyses. Amino acid counts and frequencies are computed with respect to the sequence weights.

## V. SUPPLEMENTARY MATERIAL

CCMpredPy and CCMgen are available under GNU AGPL3 at https://github.com/soedinglab/ccmgen. Code reproducing the analysis results can be found at https://github.com/soedinglab/ccmgen-scripts.

## VI. ACKNOWLEDGMENTS

This work was supported by the DFG (grant GRK 1721) and Susann Vorberg is supported by a DFG fellowship through the Graduate School of Quantitative Biosciences Munich (QBM).

